# Mycelium as a scaffold for biomineralized engineered living materials

**DOI:** 10.1101/2024.05.03.592484

**Authors:** Ethan Viles, Ethan Heyneman, Shuyi Lin, Virginia Montague, Amir Darabi, Lewis M. Cox, Adrienne Phillips, Robin Gerlach, Erika Espinosa-Ortiz, Chelsea Heveran

## Abstract

Engineered living materials (ELMs) are garnering considerable attention as a promising alternative to traditional building materials because of their potentially lower carbon footprint and additional functionalities conferred by living cells. However, biomineralized ELMs designed for load-bearing purposes are limited in their current design and usage for several reasons, including (1) low microbial viability and (2) limited control of specimen internal microarchitecture. We created ‘third generation’ biomineralized ELMs from fungal mycelium scaffolds that were mineralized either by the fungus itself or by ureolytic bacteria. Both self-mineralized (i.e. fungally-mineralized) and bacterially-mineralized scaffolds retained high microbial viability for at least four weeks in room temperature or accelerated dehydration storage conditions, without the addition of protectants against desiccation. The microscale modulus of calcium carbonate varied with the different biomineralized scaffold conditions, and moduli were largest and stiffest for bacterial biomineralization of fungal mycelium. As an example of how mycelium scaffolds can enable the design of complex internal geometries of biomineralized materials, osteonal-bone mimetic architectures were patterned from mycelium and mineralized using ureolytic bacteria. These results demonstrate the potential for mycelium scaffolds to enable new frontiers in the design of biomineralized ELMs with improved viability and structural complexity.

**Progress and Potential:** Biomineralized engineered living materials (ELMs) offer new approaches for increasing the sustainability of building materials and processes. However, the design and usage of biomineralized ELMs is constrained by several important limitations, including low microbial viability and limited ability to control internal microarchitecture. Fungal mycelium scaffolds, biomineralized by either fungi or bacteria, achieve much higher viability of ureolytic microorganisms than what has been reported for biomineralized ELMs. Further, mycelium scaffolds permit the manufacturing of complex architectures, such as inspired by the structure of osteonal bone. Mycelium scaffolds have the potential to enable new frontiers in the design and use of biomineralized ELMs.

**Graphical Abstract:** 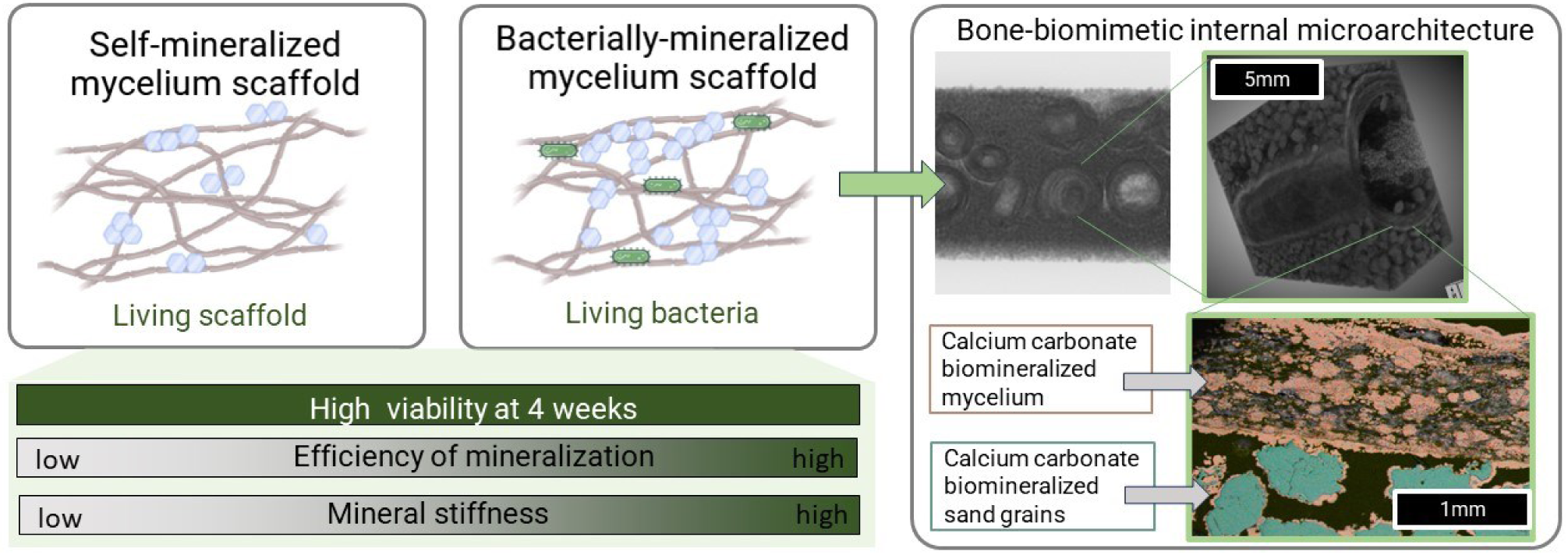

## 1. Introduction

Conventional cementitious materials that use ordinary Portland cement have an immense carbon footprint and are estimated to account for 5-8% of anthropogenic CO_2_ emissions^1^. There is strong motivation to produce building materials with lower carbon footprints, such as may be possible through leveraging microbial metabolisms to perform biocementation^2–7^. Microbial biocementation, which is achieved through microbial-induced calcium carbonate precipitation (MICP), has been used for years at the field scale to stabilize soils and seal cracks in oil wells and in recent years has been utilized to manufacture building materials appropriate for low service loads (e.g., masonry units and paving tiles)^8–10^. These types of biomineralized engineered living materials (ELMs) have seen multiple generations of development. In the ‘first generation’, MICP was used to bridge together aggregate, such as sand or crushed rock, with biomineral ^5, 8, 9, 11^. Bacterial viability was not usually the focus of the materials but was assumed to be transient. In the ‘second generation’, hydrogel scaffolds (i.e., gelatin, agarose, alginate) were used to improve the efficiency of constructing load-bearing materials, including those with complex external geometry ^4, 12–18^. Some of these materials contained living cells for a period time that conferred temporary desirable functions to the material, such as photosynthesis ^4, 16^. In this work, a ‘third generation’ of biomineralized ELMs is introduced, which improves both the cellular viability and control over internal microarchitecture through using fungal mycelium as a biological scaffold for biomineralization.

The central characteristic of these third-generation biomineralized ELMs is the use of a biological scaffold. Most mineralized materials that occur in nature such as bone, coral, and nacre are generated from the mineralization of a biological scaffold consisting of biopolymers and other proteins^19, 20^. One motivation for using a biological scaffold within ELMs is that they have the potential to be more durable than hydrogels, which degrade at ranges of humidities and temperatures common with outdoor use cases. A second motivation of using biological scaffolds is to improve microbial viability in biomineralized ELMs. In second-generation biomineralized ELMs, viability is still modest (1-4 weeks) with either special storage conditions or the inclusion of desiccant protectants^4, 16^. Importantly, these restrictions limit the amount of time a material can perform functions unique to living materials (e.g., biomineralization, photosynthesis, self-healing). A third motivation is that biopolymer scaffolds may confer control over the scaffold architecture in a manner that has not been available when working with hydrogels, which could allow complex microarchitectures of biomineralized ELMs for the first time^21, 22^.

Fungal mycelium is a promising candidate for scaffolding third-generation biomineralized ELMs. Mycelium, which contains biopolymers such as chitin, has seen a surge in use as a scaffold for soft materials, such as packaging or insulation^23–30^. Fungal mycelium is naturally resistant to heat and humidity as it exists within a variety of natural environments, and therefore, is likely to be more environmentally robust than currently used hydrogel scaffolds^26, 28^. To date, fungal mycelium has been shown to be able to achieve biomineralization in some contexts^2, 6, 31–33^. However, the use of mycelium as a scaffold for biomineralized, load-bearing materials is an emerging idea^2^. Mycelium scaffolds could also possibly harbor viable bacteria for longer amounts of time than has been reported for other types of stiff ELMs, since bacterial attachment to fungal mycelium has been extensively reported^34, 35^. Furthermore, compared to inert scaffolds, living mycelium scaffolds introduce interesting prospects for environmental responsiveness, improved self-healing, and potentially other functionalities.

Fungal mycelium could also be utilized to biomineralize the ELM. Previously, biomineralized ELMs have been stiffened by bacterial biomineralization^5, 8, 11, 12, 14, 16, 36–38^. The most common metabolism employed in these types of materials is urea hydrolysis, facilitated by the production of the urease enzyme by bacterial species such as *Sporosarcina pasteurii* ^11, 15, 39^. In the presence of urease, urea is hydrolyzed into ammonium and carbonate ions, raising the pH and alkalinity of the surrounding environment. When sufficient calcium is present calcium carbonate is precipitated. Some species of fungi, such as *Neurospora crassa*, are ureolytic and can perform fungally induced calcium carbonate precipitation (FICP) ^2, 31–33^. However, FICP is much less understood when compared with bacterially-induced calcium carbonate precipitation (BICP). FICP has been used to biomineralize sand and porous concrete, but has not been used as a scaffold for biomineralized ELMs^2, 6^. As a result, it is unclear whether self-mineralization of a fungal scaffold (via FICP) or bacterial mineralization of a fungal scaffold (via BICP) would produce more desirable material and viability attributes.

Another characteristic of mycelium scaffolds is the possibility of forming biomineralized ELMs with desirable internal architectures. Natural composite materials with desirable strength and toughness commonly have complex, optimized internal microarchitectures (e.g., bone, coral, nacre, glass sponges) ^19, 37, 38, 40–42^. Furthermore, designed internal porosity could be important for the transport of biomineralizing solutions or cellular nutrition in a scaffolded structure ^21, 22^. Structures with this degree of internal geometric complexity would be difficult to achieve using traditional hydrogel systems. The planar growth of some species of fungus, including *N. crassa*, produces a scaffold material that can be molded or shaped and then further biomineralized. One desirable internal geometry would be the osteonal structure of cortical bone. Osteons are ring-like structures with plies of biomineralized collagen, centered around an internal channel, that considerably toughen cortical bone^20, 43^ and permit nutrition/waste exchange and cellular communication ^44^.

In this work, we construct third-generation biomineralized ELMs using fungal mycelium. The mycelium scaffold was either biomineralized by itself or through biomineralizing bacteria. In either case, viability of the living component was high at 4-weeks of storage non-optimal (room temperature or elevated temperature, not humidified) storage conditions. The biomineralization efficiency as well as the stiffness of the biomineral were greatest for bacteria-mineralized fungal scaffolds. These results, together with our demonstration of osteonal-mimetic internal microarchitectures, demonstrate the potential of fungal mycelium to overcome several limitations of previous generations of biomineralized ELMs.

## 2. Results

### 2.1 Mycelium scaffolds can self-mineralize but efficiency depends on nitrogen source

To investigate how *Neurospora crassa* mycelium self-mineralizes, batch studies were performed over 10 days where *N. crassa* was cultured with growth media, urea, and calcium. These batch studies demonstrate that *Neurospora crassa* can perform fungally induced calcium carbonate precipitation in malt media (FICP-malt) with some decrease in solution urea and calcium (**Figure 1A-B**, **Table 1**). In an attempt to increase fungal ureolysis, *N. crassa* scaffolds were cultured in a defined medium where urea was the sole source of nitrogen (FICP-def). FICP-def scaffolds utilized slightly more urea and twice the amount of calcium over 10 days than the FICP-malt scaffolds (**Figure 1A-B**, **Table 1**). Thus, we determine that FICP-def is more efficient at biomineralization than FICP-malt. Biomineral accumulation on the fungal scaffolds, estimated by mass lost after an acid digest, was 20% greater for FICP-def than FICP-malt (**Figure 1D**). Both FICP-def and FICP-malt fungal scaffolds had a higher biomineral fraction than the control of abiotically mineralized mycelium scaffold (AICP, **Table 1**). The greater biomineral mass fraction over the same time (10 days) is interpreted as improved biomineralization efficiency of the FICP-def condition (**Figure 1**, **Table 1**).

**Figure 1:**
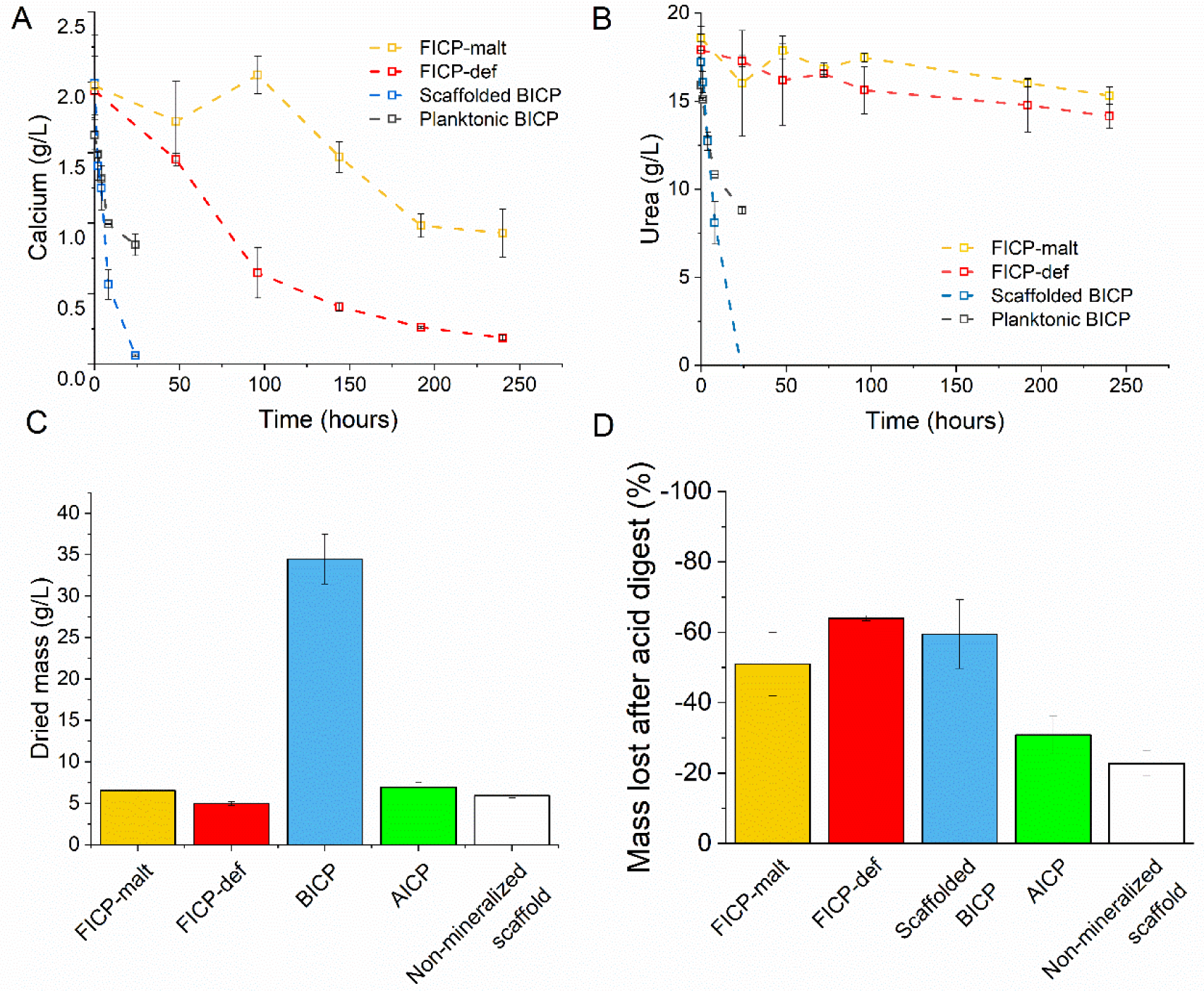
Self- or bacteria-mineralization of fungal mycelium. A) Dissolved calcium concentrations and B) dissolved urea concentrations during biomineralization for FICP-malt and FICP-def of living fungal mycelium, planktonic BICP (no mycelium), AICP (abiotic mineralization) of autoclaved mycelium, and scaffolded BICP of autoclaved mycelium. Points are means of measurements from biological triplicates with error bars showing the standard deviation. C) Constant mass of dried biomineralized scaffolds after mineralization and non-mineralized mycelium after 10 days of growth. Bars are the means of measurements from biological triplicates with error bars showing the standard deviation. D) Percent mass loss after nitric acid digest normalized to scaffold starting dry weight to estimate calcium carbonate content of scaffolds. Bars are mean measurements from biological triplicates with error bars showing the standard deviation.

**Table 1:**
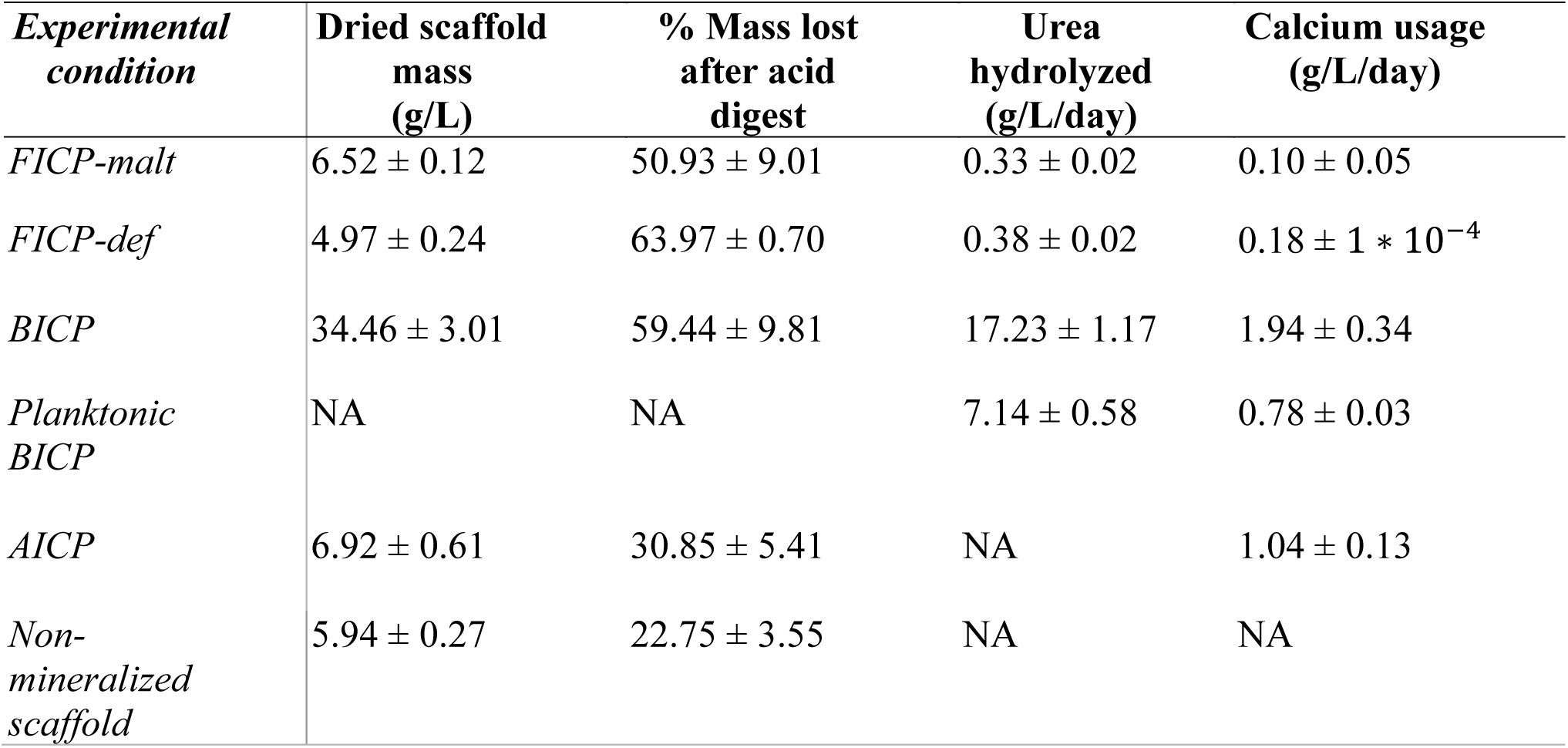
Biomineralization efficiency of scaffolded ELMs and non-scaffolded controls.

| **Experimental condition** | **Dried scaffold mass (g/L)** | **% Mass lost after acid digest** | **Urea hydrolyzed (g/L/day)** | **Calcium usage (g/L/day)** |
| --- | --- | --- | --- | --- |
| *FICP-malt* | 6.52 ± 0.12 | 50.93 ± 9.01 | 0.33 ± 0.02 | 0.10 ± 0.05 |
| *FICP-def* | 4.97 ± 0.24 | 63.97 ± 0.70 | 0.38 ± 0.02 | 0.18 ± 1 * 10<sup>-4</sup> |
| *BICP* | 34.46 ± 3.01 | 59.44 ± 9.81 | 17.23 ± 1.17 | 1.94 ± 0.34 |
| *Planktonic BICP* | NA | NA | 7.14 ± 0.58 | 0.78 ± 0.03 |
| *AICP* | 6.92 ± 0.61 | 30.85 ± 5.41 | NA | 1.04 ± 0.13 |
| *Non-mineralized scaffold* | 5.94 ± 0.27 | 22.75 ± 3.55 | NA | NA |

### 2.2 Mycelium scaffolds are efficiently biomineralized by ureolytic bacteria

To the best of our knowledge, the ability of *S. pasteurii* to biomineralize mycelium scaffolds has not yet been studied. Here, we investigated the ability of *S. pasteurii* to colonize and biomineralize non-viable *N. crassa* scaffolds. Mycelia grown in non-mineralizing medium were autoclaved prior to being added to cultures containing *S. pasteurii* to assess the potential of bacterial colonization and biomineralization of the scaffolds. After the incubation period, fungal scaffolds were analyzed for bacterial colonization with SEM and confocal microscopy. In the mineralized samples, SEM images show minerals along the fungal hyphae accompanied by the presence of bacteria. In non-mineralized scaffolds, the presence of bacteria is less evident. (**Figure 2A-B**). Confocal laser scanning microscopy (CLSM) of non-mineralized scaffolds also suggest decreased presence of bacteria (**Figure 2C-E**).

**Figure 2:**
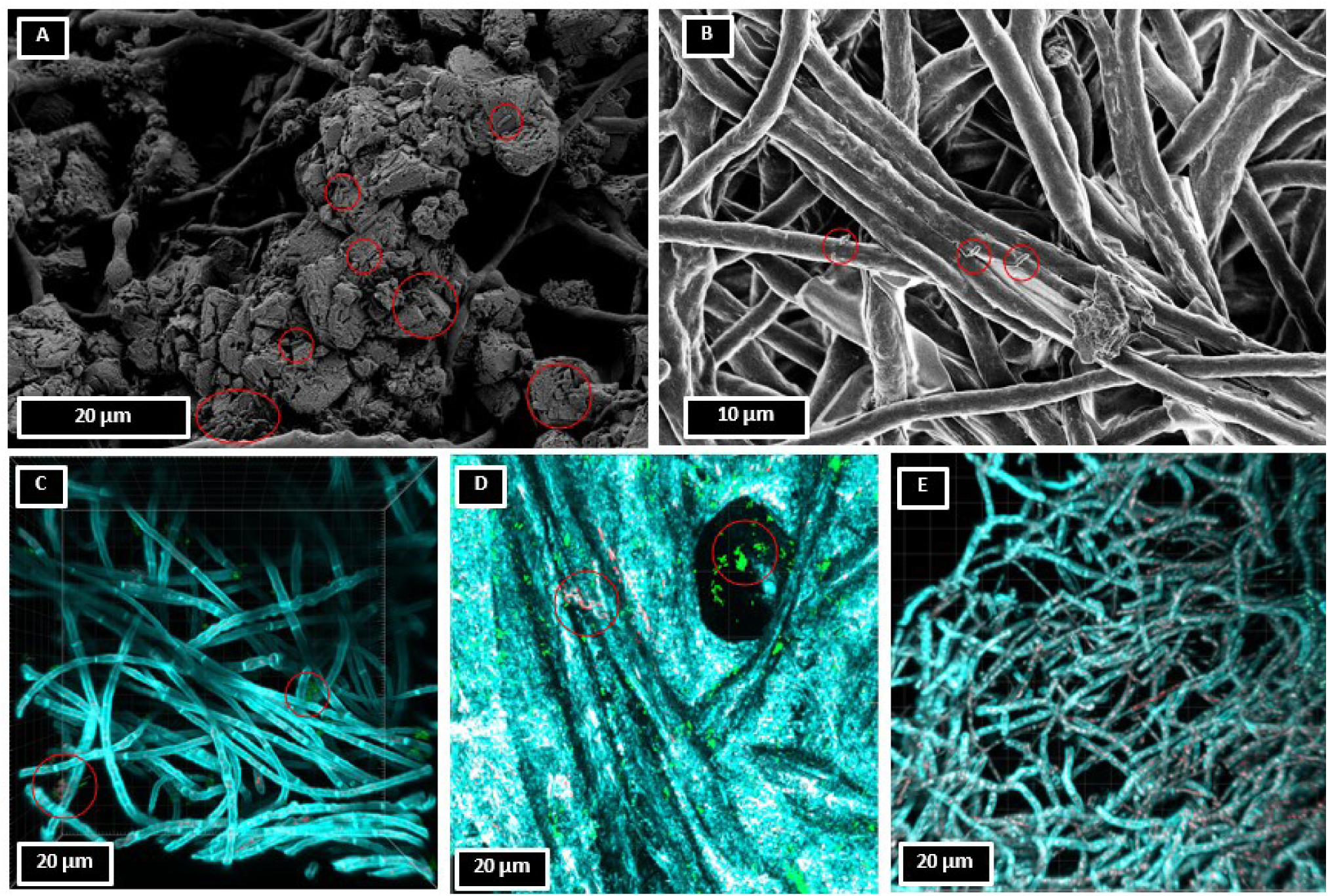
SEM and confocal microscopy images of S. pasteurii growing in liquid cultures with non-viable mycelium scaffolds. Red circles indicate the presence of bacteria. **A**) SEM image of non-viable mycelium scaffolds mineralized by *S. pasteurii*. **B**) SEM image of non-viable mycelium scaffolds (non-mineralized) inoculated with *S. pasteurii.* **C**) CLSM image of non-viable mycelium scaffolds inoculated with *S. pasteurii* but not mineralized. Green indicates viable bacteria, red indicates membrane compromised bacteria, and cyan indicates fungal mycelium. **D)** CLSM image of a cellulose coupon with only *S. pasteurii*. Green and red indicate viable and membrane-compromised bacteria, respectively. The cyan color is from the background coloration of the coupon. **E)** CLSM image of *N. crassa* mycelium only. The fungal mycelium stains cyan and the red indicates either some uptake of propidium iodide or autofluorescence of fungal cells.

To compare *N. crassa* self-mineralization to bacterial mineralization, the ability of *S. pasteurii* to induce calcium carbonate precipitation (BICP) on autoclaved *N. crassa* scaffolds was tested. Bacterial biomineralization was more efficient than both self-mineralization methods (FICP-malt and FICP-def) as it required just 24 hours to hydrolyze all urea and to remove all dissolved calcium present in solution (**Figure 1A-B**, **Table 1**). Acid digestion of the scaffolds estimates that calcium carbonate made up about 50% of the scaffolds’ weight (**Figure 1D**, **Table 1**). The BICP and FICP-def scaffolds had a similar percent mass of calcium carbonate accumulation (60% and 65%, respectively). Because mycelium scaffolds used for bacterial mineralization were first grown on malt media with no urea or calcium, the overall scaffold mass was much greater for BICP than for fungal self-mineralization (i.e. FICP-malt and FICP-def) conditions as demonstrated by dried mass measurements (**Figure 1C**, **Table 1**).

### 2.3 Living, ureolytic components of biomineralized fungal scaffolds maintain viability for at least 4 weeks

For self-mineralized (FICP-malt or FICP-def) scaffolds, the *N. crassa* scaffold is the living component. In BICP scaffolds, *S. pasteurii* is the living component and the *N. crassa* scaffold is non-viable. The viability of each type of living component was determined after biomineralization and drying for up to four weeks at both room temperature (23°C ± 1°C) and 30°C (**Table 2**). Self-mineralized mycelium (FICP-def and FICP-malt) scaffolds demonstrated growth when introduced to fresh media at four weeks following drying at either room temperature or 30°C (**Figure 3**, **Table 2**). Mechanically disturbing the mineralized scaffolds was required for some samples to instigate this growth (**Figure 3A**). Non-mineralized mycelium also retained viability at 4 weeks after drying at either temperature but did not require mechanical disturbance of any samples to result in the growth of mycelium (**Figure 3**).

**Figure 3:**
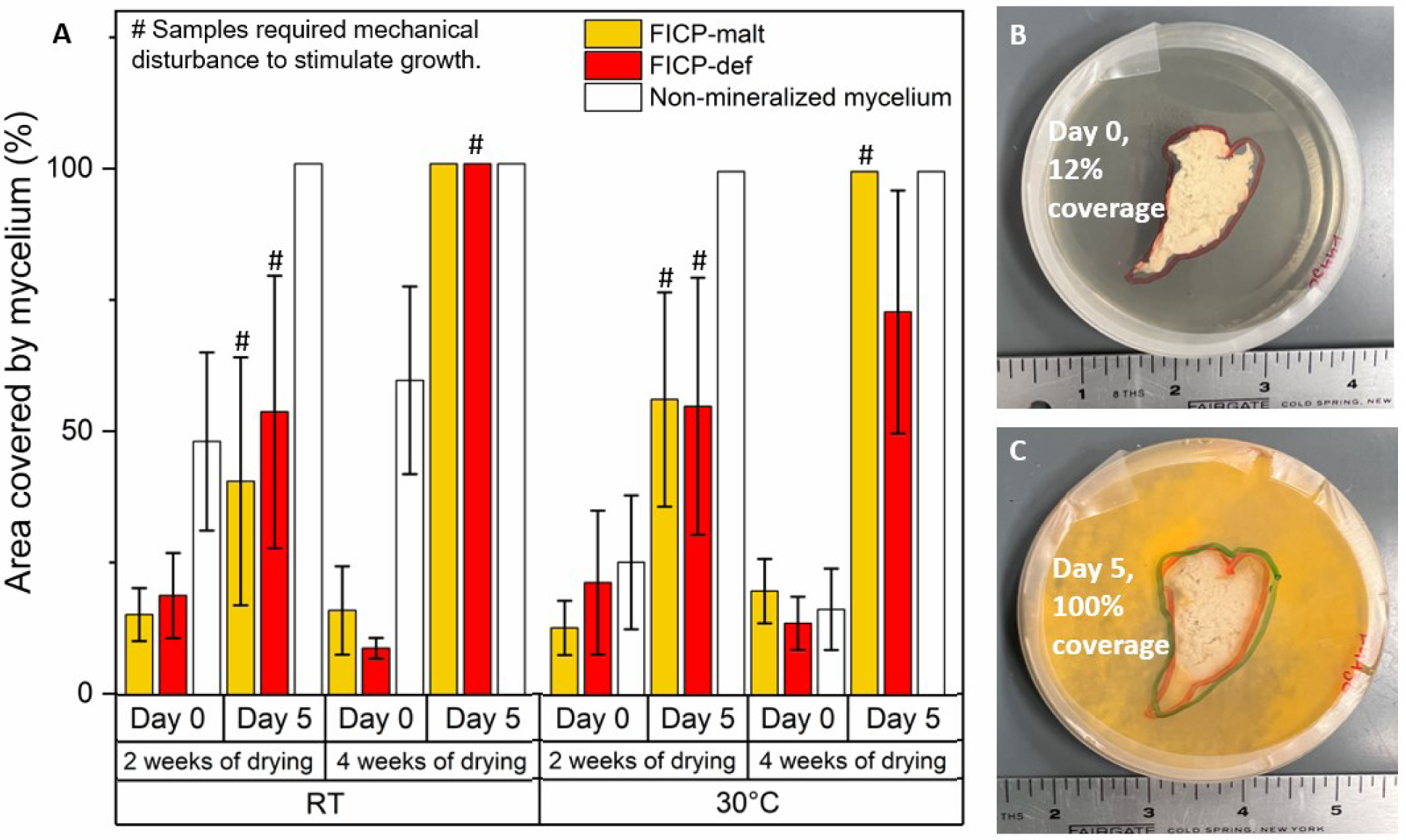
Growth of *N. crassa* after mineralization and drying. A) Growth of *N. crassa* after drying at room temperature (RT) or 30°C. Growth was measured after two weeks and four weeks of drying. Growth was measured by calculating percent area covered on agar plates during incubation. Growth measurements were made at day 0 and day 5 of incubation. Each bar represents a mean measured from biological triplicates and errors bars are the standard deviation for those measurements. Some measurements show no standard deviation because each plate had 100% coverage. B) Day 0 of inoculating mycelium scaffolds (FICP-malt) that were dried for four weeks at 30°C. C) Day 5 of growth from the same scaffold shown in Figure 3B. Each bar represents a mean measured from biological triplicates and errors bars are the standard deviation for that measurement. Some measurements show no standard deviation because each plate had 100% coverage.

**Table 2:** Biomineralized scaffolds show viability of scaffold or bacteria living components for 4 weeks.

| *Scaffold Type* | **Abbreviation** | **Mineralizing organism** | **2-week viability** | **4-week viability** |
| --- | --- | --- | --- | --- |
| *Self-mineralized fungal scaffold, malt medium* | FICP-malt | *N. crassa* | Viable | Viable |
| *Self-mineralized fungal scaffold, defined medium* | FICP-def | *N. crassa* | Viable | Viable |
| *Bacterially-mineralized fungal scaffold* | BICP | *S. pasteurii* | Viable | Viable |

Non-viable mycelium scaffolds inoculated with *S. pasteurii* retained high amounts of viable bacteria after mineralization and drying. Following *S. pasteurii* biomineralization of *N. crassa* scaffolds, these scaffolds contain an abundance of culturable *S. pasteurii* cells (>1×10^4^ CFU/mL) after both 2 and 4 weeks of drying at either room temperature or 30°C (**Figure 4A**). In contrast, no colonies were recovered from planktonic *S. pasteurii* cultures after drying in an oven at 30°C for 4 weeks (**Figure 4A**). This indicates that the fungal scaffolds might increase the resistance of *S. pasteurii* cells against temperature or desiccation stress. Since the CFU plate counts are not specific to *S. pasteurii* growth, the presence of *S. pasteurii* was reconfirmed by measuring the ability of surviving cells to hydrolyze urea. Urea hydrolysis was achieved at every combination of temperature and drying time for *S. pasteurii* associated with mycelium scaffolds. However, strikingly, the extent of urea hydrolysis was similar between freshly prepared BICP scaffolds and BICP scaffolds dried at room temperature for 4-weeks, for example, 2 g/L of urea left after 24 hours of incubation (**Figure 4B**). The extent of ureolysis was much greater for scaffolded samples dried at room temperature than for scaffolded samples dried at 30°C, and greater than planktonic samples across the board.

**Figure 4:**
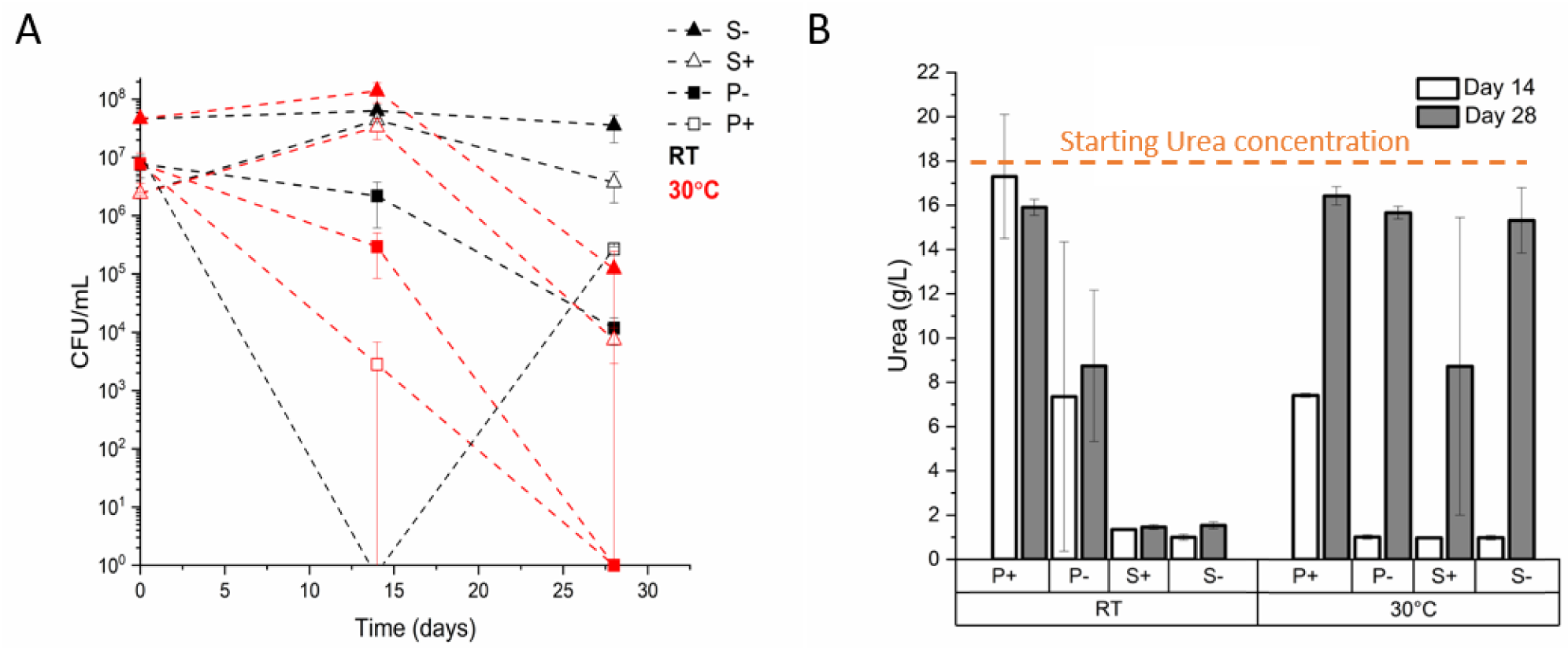
Bacterial growth and ureolytic activity of mineralized bacteria (+), non-mineralized bacteria (-), bacteria grown on a mycelium scaffold (S), and bacteria grown planktonically, without a scaffold, (P) after drying for four weeks. A) Plate counts of *S. pasteurii* cultured for 24 hours after drying at 30°C or room temperature (RT) for four weeks. Each point represents a mean measured from biological triplicates, and each error bar is the standard deviation for those measurements. Red indicates samples dried at 30°C while black indicates samples dried at room temperature (RT). B) Ureolytic activity of *S. pasteurii* cultured for 24 hours after drying at 30°C or room temperature (RT) for four weeks. Each bar represents a mean measured from biological triplicates and errors bars are the standard deviations of those measurements.

### 2.4 Biomineral characteristics and morphology differ between self-biomineralized and bacteria-biomineralized fungal scaffolds

Because the size, morphology, and stiffness of biomineral would be expected to impact the utility of biomineralized scaffolds as load-bearing materials, these characteristics were evaluated using several imaging and indentation techniques.

The morphology and location of biominerals differed with the type of biomineralization method. Scanning electron microscopy with electron-dispersive spectroscopy (SEM-EDS) demonstrates that BICP scaffolds have large calcium-containing biominerals that bridge individual hyphae (**Figure 5-6**). By contrast, FICP-malt and AICP scaffolds have calcium-containing minerals deposited predominately onto the top of the mycelium mat. FICP-def scaffolds show hyphae encrusted with small calcium-containing biominerals that do not bridge the hyphae together (**Figure 5-6**).

**Figure 5:**
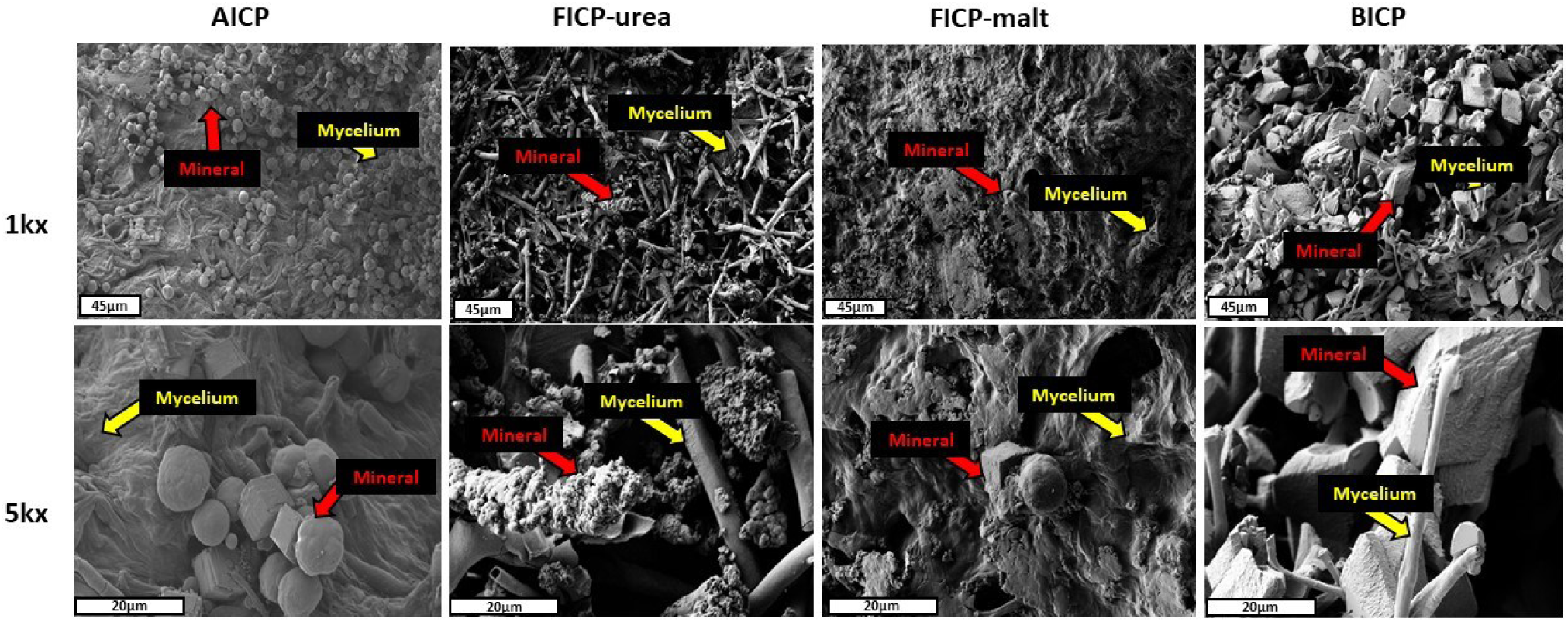
Self-mineralized, abiotically mineralized, and bacterially-biomineralized scaffolds. Column 1 shows images from self-mineralized fungal induced calcium carbonate precipitation FICP living mycelium samples using malt broth extract FICP-malt. Column 2 shows images from FICP living mycelium where the only source of nitrogen was urea in the medium FICP-def. Column 3 shows images from autoclaved mycelium scaffolds mineralized with abiotically induced calcium carbonate precipitation AICP. Column 4 shows images from autoclaved mycelium scaffolds mineralized utilizing the bacterium *S. pasteurii* BICP. Rows within this image correspond to the magnification of each image 1kx (1000x) and 5kx (5000x).

**Figure 6:**
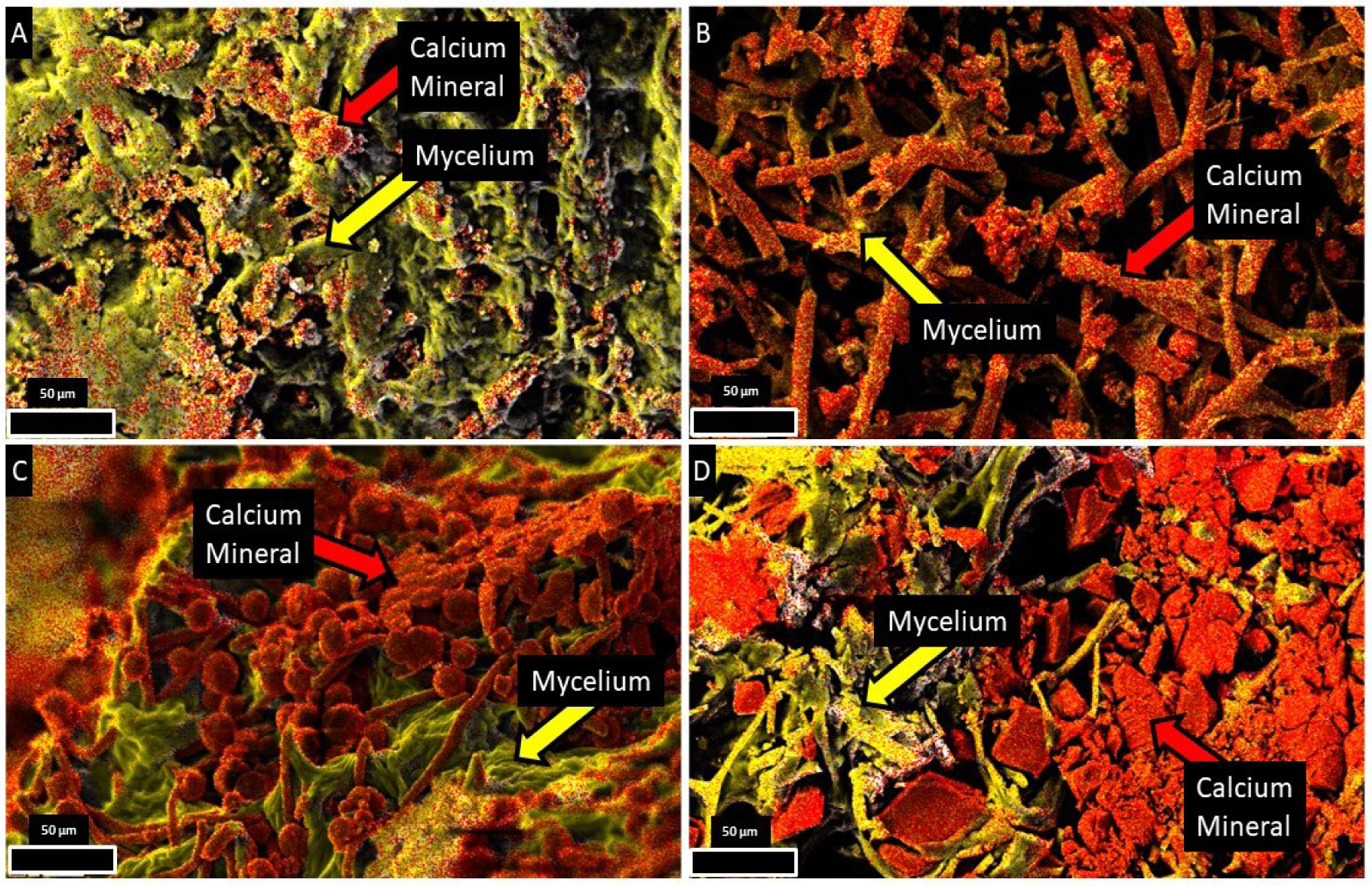
SEM-EDS images of mineralized mycelium from autogenous and exogenous mineralization. Scaffolds include: A) FICP-malt living mycelium, B) FICP-def living mycelium, C) AICP autoclaved mycelium, D) BICP autoclaved mycelium. Carbon is labeled yellow, and calcium is labeled red in all images.

XRD confirmed that the calcium minerals identified on BICP, FICP-malt, FICP-def, and AICP scaffolds by SEM-EDS are indeed calcium carbonate (**Figure 7**). All mineralized scaffolds showed calcium carbonate as the only calcium mineral present, and every scaffold contained calcite and vaterite polymorphs of calcium carbonate. XRD data also indicate that the ratio of vaterite and calcite might vary between mineralized scaffolds.

**Figure 7:**
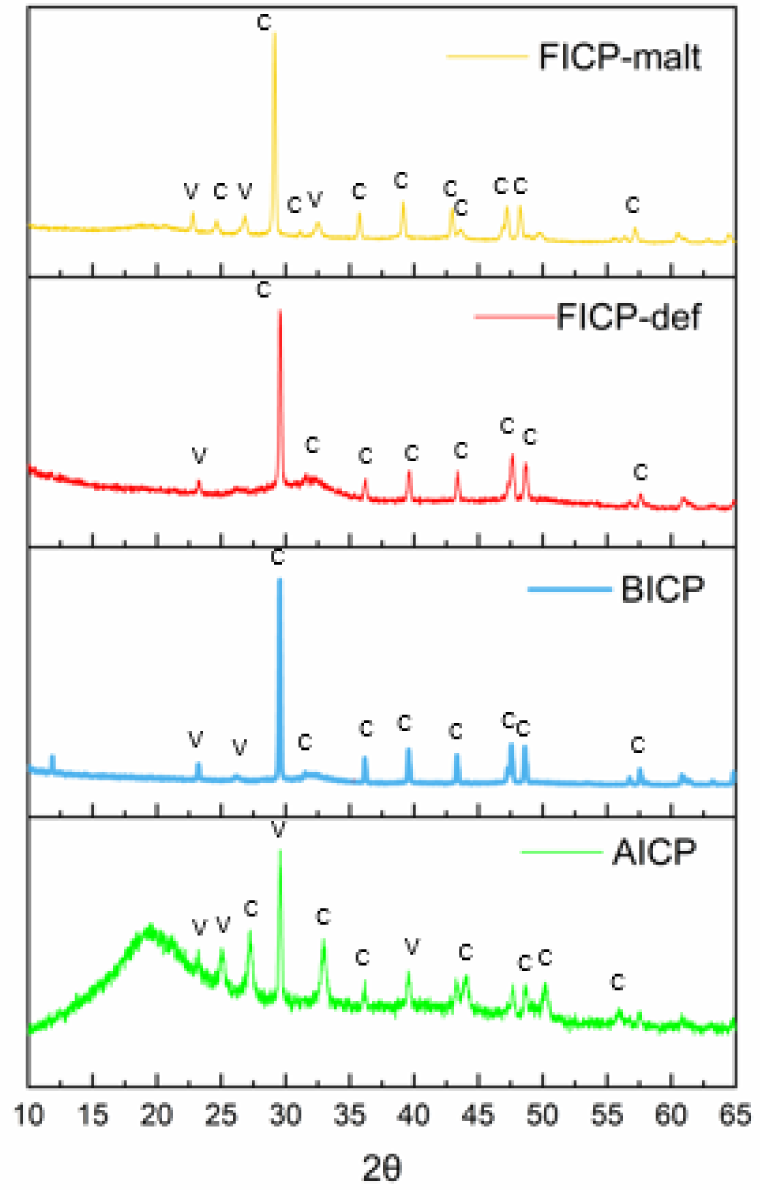
XRD identification of minerals precipitated on fungal mycelium scaffolds. FICP-malt, FICP-def, BICP, and AICP scaffolds all show calcite (‘c’) and vaterite (‘v’) precipitation.

Calcium carbonate moduli also varied with the biomineralization condition (**Figure 8A**). As evaluated at the microscale by instrumented nanoindentation, FICP-def scaffolds had 276% stiffer mineral than FICP-malt. BICP scaffolds had 632% and 230% stiffer mineral than FICP-malt and -def scaffolds, respectively.

**Figure 8:**
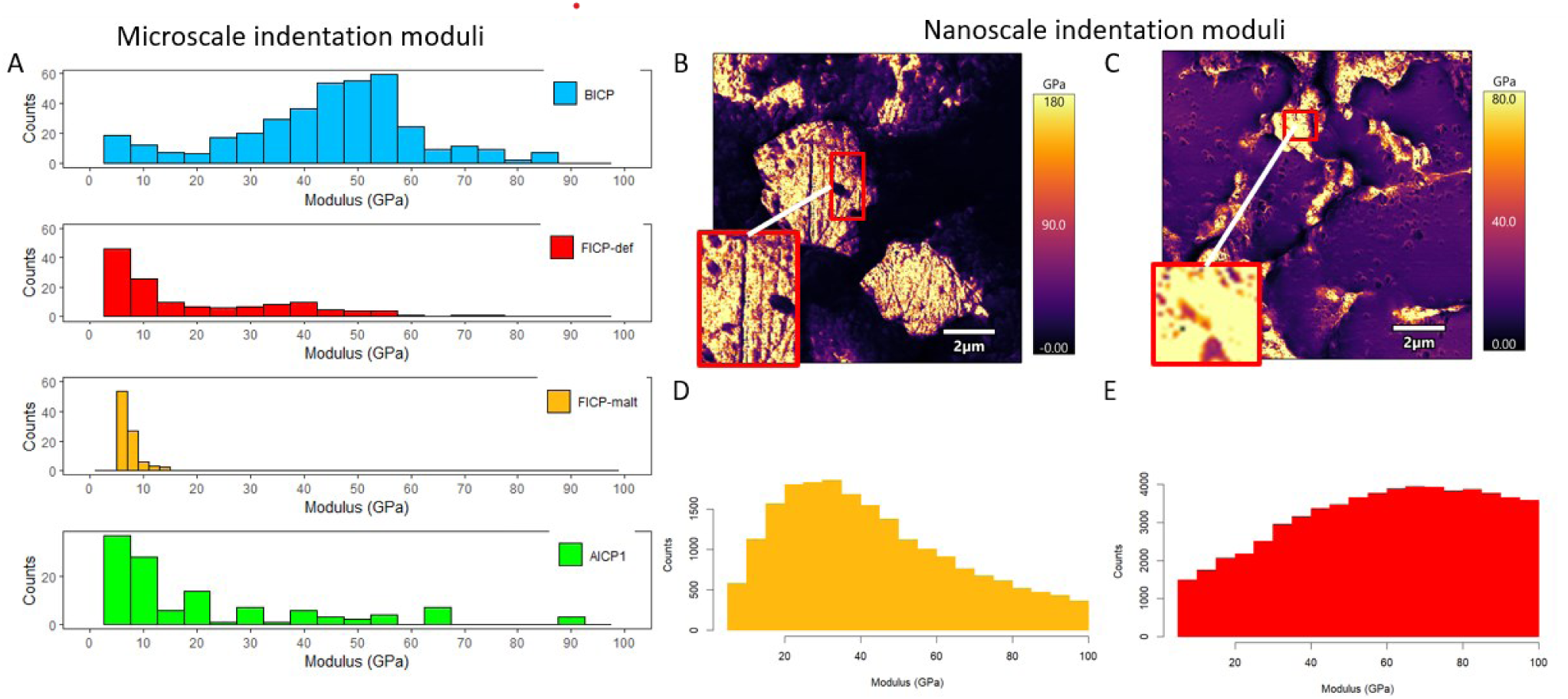
Biomineral moduli depend on mineralization condition. A) Histograms of nanoindentation moduli measured by microscale nanoindentation. Each histogram represents a different mineralization condition: blue is BICP, red is FICP-def, yellow is FICP-malt, and green is AICP. The data were acquired from grids at a random location that had sufficient mineral for indentation. B) A representative map of AFM moduli from an FICP-malt scaffold. C) A representative map of AFM moduli from an FICP-def scaffold. D) A histogram of FICP-malt moduli measured with atomic force microscopy (AFM). E) A histogram of FICP-def moduli measured with AFM.

The low biomineral moduli for the FICP-malt condition could potentially be attributed to either low stiffness of calcium carbonate, inclusion of organics within the mineral, or a composite response to nanoindentation (tip contact area ∼ 2-3 µm^2^) between the small calcium carbonate minerals and the mycelium and epoxy embedding material. To supplement these microscale indentation measurements, atomic force microscopy (AFM) was employed to map calcium carbonate stiffness at a higher resolution (∼40 nm). AFM mapping demonstrated that the modulus for FICP-malt samples, evaluated for calcium carbonate biominerals away from interfaces with epoxy and small pores, is indeed lower than FICP-def samples (**Figure 8D-E**). For both tested conditions, moduli calculated from AFM force curves exhibit larger moduli than microscale indentation measurements. These data demonstrate that porosity and/or epoxy intrusion likely influence the modulus measured at the microscale.

### 2.5 Osteonal-mimetic microarchitecture within biomineralized ELMs

Current biomineralized ELM scaffolds do not permit control over internal microarchitecture, but the material properties of biomineralized natural composites often benefit from complex internal porosity or interfaces^19, 22, 38, 40, 44^. With the usage of bacterially-mineralized fungal scaffolds (BICP) in this study, we had an opportunity to control the internal microarchitecture of biomineralized ELMs for the first time. We constructed biomineralized ELMs with an internal microarchitecture inspired by cortical bone. Specifically, we employed a design inspired by osteons, which are ring-like structures of concentric lamellae found within the cortical bone of large animals that serve to both vascularize and toughen bone tissue^45, 46^. To form these osteon-inspired structures internal to a larger composite, autoclaved sheets of fungal mycelia were bacterially-mineralized to form columns of concentric mineralized bands (**Figure 9B-C**). These were then placed in a prism-shaped mold (of 2.54 x 2.54 x 10.16cm) with voids filled with sand and mineralized again with *S. pasteurii* (**Figure 9A**). SEM and SEM-EDS demonstrate successful biomineralization throughout the osteon-inspired structures. SEM-EDS and microCT also demonstrate that the biomineralized osteon-inspired structures have clear concentric rings of biomineralized scaffold that are easily identified from sand particles (**Figure 9A-C**).

**Figure 9:**
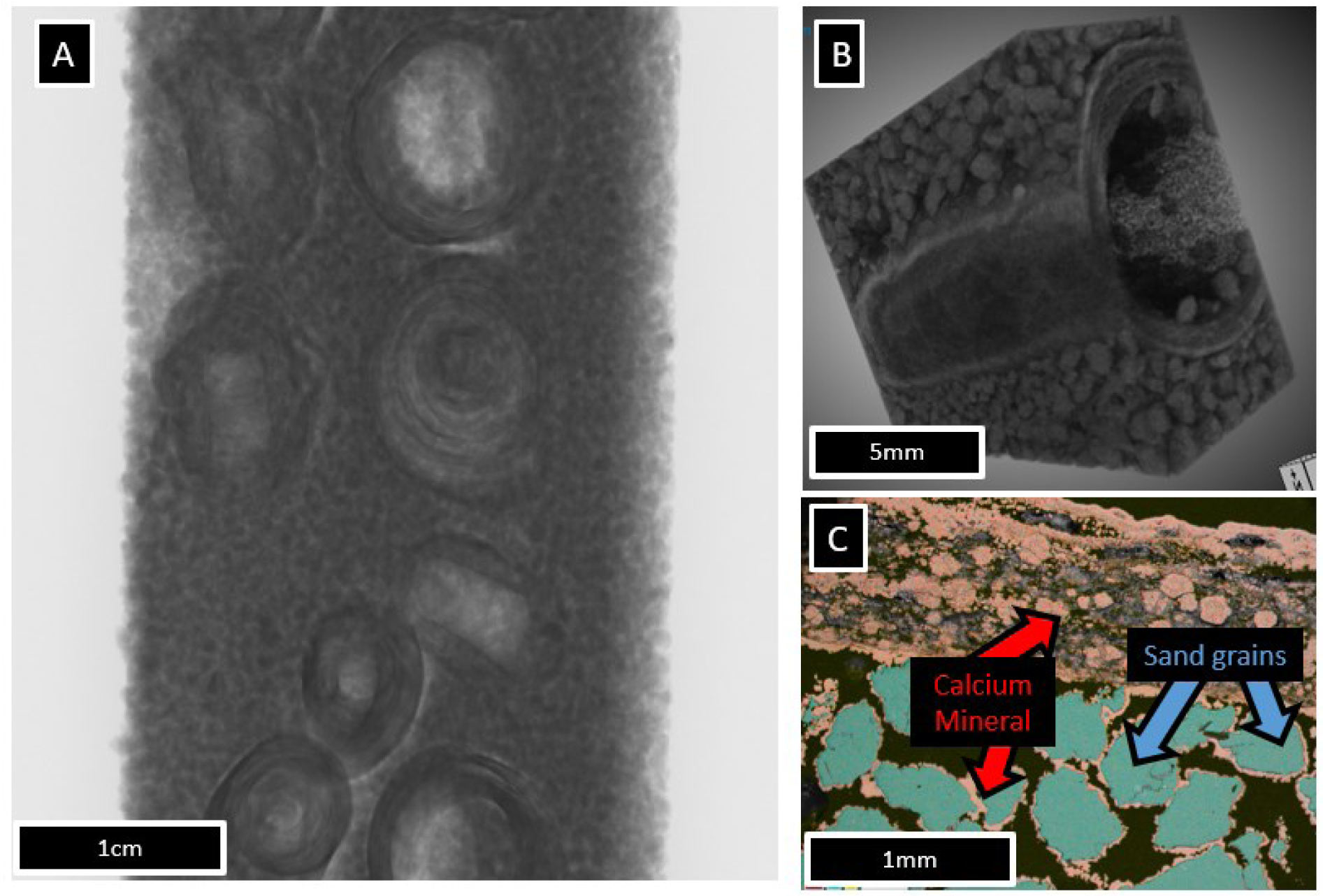
Mycelium scaffolds can be used to design interior architectures of biomineralized prisms. A) MicroCT scan of osteon-mimetic architectures within a biomineralized prism. B) MicroCT imagery of a single artificial osteon. C) SEM-EDS imagery of the artificial osteon prism. Blue indicates silica, while red and yellow indicate calcium and carbon, respectively. The SEM image used is from a backscatter electron detector, and the brighter pixels indicate elements with higher atomic mass.

## 3. Discussion

Biomineralized ELMs (sometimes called ‘living building materials’, LBMs) are attractive for building and infrastructure material applications because of their potential for more sustainable manufacturing compared to cementitious materials ^4, 5, 15, 16, 21^. Further, living cells within ELMs can enable additional desirable functionalities to these materials, such as regenerative properties, environmental responsiveness, or self-healing ^5, 13, 15, 16, 47–51^. Much progress has been made in first- (i.e., non-scaffolded) and second-generation (i.e., hydrogel-scaffolded) biomineralized ELMs and some of these materials are now entering commercial usage ^4, 15, 16, 21, 37, 47, 51, 52^. However, there are important limitations to these technologies that should be surmounted to improve when and how biomineralization can be beneficially utilized. First, the microbial viability within these materials is limited to just a few days or weeks ^4, 7, 16, 21^. Second, hydrogel scaffolds have limitations in their environmental durability that decrease their appropriateness for many outdoor applications ^53, 54^. Third, the internal geometry of these materials is difficult to control, which limits the construction of many bio-inspired or otherwise geometrically optimized designs that may benefit material strength or cellular viability. Here, we introduce a third-generation of biomineralized ELMs that are scaffolded using fungal mycelium, with or without the addition of ureolytic bacteria.

Fungal mycelium has become popular as a scaffold for soft materials but is much less commonly used in applications involving biomineralization ^23, 24, 28–30, 55–58^. *N. crassa*, used in our study, is an example of a ureolytic fungus that has the potential for accomplishing biomineralization via urea hydrolysis. We found that *N. crassa* can achieve self-mineralization and that the efficiency of this biomineralization was influenced by media nitrogen availability. When nitrogen in media was limited to urea, more biomineralization was witnessed. By contrast, a nitrogen-rich fungal medium promoted more scaffold growth but less mineral formation. However, the most efficient strategy to promote robust scaffold biomineralization was to grow the fungus in rich, non-mineralizing media and then biomineralize the grown, autoclaved scaffold using highly efficient ureolytic bacteria (i.e., *S. pasteurii*). These results are likely influenced by different amounts of ureolytic biomass in each condition.

The characteristics of calcium carbonate produced through fungal scaffold biomineralization also depended on the manner of biomineralization. The stiffest biominerals were produced by bacterial biomineralization. These calcium carbonate minerals had similarly stiff crystals that are reported in various other works where bacteria biomineralize other scaffolds red is FICP-def^12, 59^. Further, the bacterial-produced calcium carbonate effectively interpenetrated the fungal hyphae network. These characteristics are desirable for producing a strong scaffold ^9^. Fungal self-mineralization produced biominerals with varying stiffnesses that were often much less stiff than bacterially produced biomineral. FICP-malt produced much smaller and less-stiff biominerals than FICP-def. It is possible that the high organic content of the rich media influenced these results. In other work, calcium carbonate precipitated in the presence of high organic content produces smaller and much less stiff crystals ^11, 36^. Our work adds to a growing body of research demonstrating that calcium carbonate produced in association with ureolytic microorganisms or enzymes varies considerably in morphology and material properties and extends these findings into the fungal kingdom ^2, 4, 8, 9, 12, 15, 21, 37, 39^. These data also suggest that there is a large design space available for tailoring the growth and biomineral characteristics of biomineralized fungal scaffolds.

Biomineralized fungal scaffolds achieved much greater viability than has been reported for earlier studies of first- and second-generation biomineralized ELMs. Living fungal mycelium scaffolds (FICP-def and FICP-malt) showed high viability after biomineralization, whether drying at room temperature or slightly elevated temperature conditions (30°C) for four-weeks. In some cases, physically crushing the biomineralized mycelium and introducing nutrition appeared to revive the fungus to a metabolically active condition. *S. pasteurii* on fungal scaffolds showed very high viability (i.e., as high, or higher, than the bacterial counts taken before drying) and continued ureolytic activity after 4 weeks of storage at room temperature. Viability was also high after 4 weeks of elevated temperature exposure (30°C), but ureolytic activity was decreased.

Viability was not tested beyond 4 weeks, but it is likely substantially longer for some of our experimental conditions, based on the high prevalence of culturable, ureolytic organisms at that time point. The viability of living fungal scaffolds or ureolytic bacteria supported by these scaffolds are much greater than what has been achieved by most previous studies of microbes in either concrete or biomineralized ELMs. In cement and concrete, microbial viability is limited to few days, at best, in most instances^1, 5^. Our results also improve on most reports from hydrogel-sand systems, where microbial viability can be limited to 1-2 weeks unless special storage conditions or anti-desiccants are provided ^4, 13, 14, 16, 47–51, 60^. This improved viability enables the possibility of self-healing or environmental responsiveness for longer than has been possible for investigations of ELMs for building material applications to date. Importantly, bacterially colonized scaffolds retained more viable cells than planktonic bacterial cultures, mineralized or not. Our work does not definitively establish why the scaffolds retain higher viability. It is possible that the scaffold itself preserves sufficient moisture to aid microbial survival. It is also possible that the autoclaved mycelium scaffolds make available carbon and nitrogen that support bacterial survival. We did not explore these possibilities in the current investigation, but they would benefit from future investigation.

We also demonstrated that mycelium scaffolds can permit the construction of specimens with complex internal microarchitectures. Osteonal bone was chosen as a demonstration since this microarchitecture is well-known to contribute to bone’s excellent properties while provide important physiological functions ^20, 22, 24, 43, 44^. Mycelium scaffolds were bacterially-biomineralized into concentric rings that mimic the lamellae within osteons, and then were mineralized together with a dispersed sand phase within a beam-shaped mold. In some of these scaffolds, additional mycelium was included between the sand, which better mimics interstitial bone between osteons. Imaging studies demonstrate that the osteon-inspired mycelium layers were biomineralized, while the central canals were usually not, which may be attractive for the design of ‘vasculature’ within biomineralized ELMs. These results point towards a much larger design space for the microarchitecture of biomineralized ELMs than has previously existed.

There were several important limitations to this study. Co-cultures of living *N. crassa* and *S. pasteurii* were not viable in this investigation, which then required that *N. crassa* scaffolds were autoclaved before inoculation with *S. pasteurii*. The reasons for this lack of compatibility have not yet been determined. While we succeeded at creating biomineralized specimens of testable size and shape (e.g., of 2.54 x 2.54 x 10.16cm), we have not yet characterized their mechanical properties. Optimizing internal microarchitecture may influence the properties of these materials, but this has not yet been tested. Our initial progress in identifying fungal mycelium as a promising biological scaffold for biomineralized ELMs lays the groundwork for these next investigations.

This work establishes the potential to utilize fungal scaffolds for the manufacturing of third-generation, biomineralized ELMs. Both types of fungal scaffolds (e.g., self-mineralized living fungal scaffolds or bacteria-mineralized killed fungal scaffolds) have potential as useful ELMs and show excellent viability characteristics of the living component. We anticipate that the improved viability characteristics of these scaffolds, together with the formability of fungal mycelium into useful shapes that confer control over internal microarchitecture, will enable the design of new materials that are able to surmount critical design limitations of earlier generations of ELMs designed for load-bearing applications.

## 5. Materials and Methods

### 5.1 Growth and Mineralization Media

Several mineralization and growth media were used throughout these experiments for either bacterial growth, fungal growth, bacterial mineralization, or fungal mineralization. All growth media were prepared using deionized (DI) water. Media not containing urea was sterilized by autoclaving. Since autoclaving may degrade urea, media containing urea were filter sterilized using 0.2µm sterile filters (ThermoFisher, 595-4520). pH adjustments were made using 40 g/L NaOH and/or 36.45 g/L HCl.

*Neurospora crassa* was cultured on 15 g/L of Malt Broth Extract (MBE, Research Products International M22500) with the pH adjusted to 6.5. Fungal biomineralization was performed with two different media: MBE media that was supplemented with 20 g/L urea and 7.35 g/L of CaCl_2_ · 2H_2_O with the pH adjusted to 6.5, and a defined fungal media with urea as the only nitrogen source (FICP-def), which consisted of 5 g/L glucose, 20 g/L urea, 7.35 g/L of CaCl_2_, 0.5 g/L MgSO_4_ · 7H_2_O, 2 g/L KH_2_PO_4_ and 5 mL of trace element solution, with the pH adjusted to 4.3. Trace element solution consisted of 3 g/L MgSO_4_ · 7H_2_O, 0.5 g/L MnSO_4_, 1 g/L NaCl, 0.1 g/L FeSO_4_· 7H_2_O, 0.01 g/L IKSO_42_ · 12H_2_O, 0.01 g/L Na_2_MoO_4_ · 2H_2_O, 1.5 g/L nitrilotriacetic acid, 0.1 g/L ZnSO_4_ · 7H_2_O, 0.1 g/L CuSO_4_ · 5H_2_O, and 0.01 g/L H_3_BO_3_ ^61^.

*Sporoscarcina pasteurii* was cultured on 37 g/L of brain heart infusion (BHI, Difco 211059) supplemented with 20 g/L urea (BHI+), and the pH was adjusted to 6.5. BICP mineralization of fungal scaffolds used two media, a calcium mineralization medium (CMM+) and a calcium free medium (CMM-). CMM-consisted of 3 g/L nutrient broth (Difco 234000) and 10 g/L NH_4_Cl. CMM+ consisted of 3 g/L nutrient broth and 10 g/L NH_4_Cl, and 7.35 g/L of CaCl_2_ · 2H_2_O.

### 5.2 Microorganisms

*N. crassa* (Fungal Genetics Stock Center #2489) cultures were maintained on MBE agar plates and incubated for three days at 30°C before storage at 4°C. *N. crassa* inoculums were started by placing an agar plug with a diameter of 1 cm from plate cultures into 100 mL of MBE media. The fungus was grown in triplicate for three days on an orbital shaker table (New Brunswick Scientific, G24) at 30°C and 150RPM, covered from light sources. A 3-day-old fungal culture was then homogenized using a mechanical homogenizer (IKA T25 Digital Ultra Turrax). After homogenization, the fungal biomass was centrifuged (6000 RPM, 10 min, 4°C) and the supernatant was removed. The fungal pellet was washed with 10 mL of phosphate buffered saline (PBS) and centrifuged (6000 RPM, 10 min, 4°C). The supernatant was removed, and the wash step was repeated once more. The final volume of the homogenized fungal biomass was adjusted to 35 mL, and 1 mL aliquots of this was used to inoculate experimental samples. The dry weight of the two remaining 3-day old fungal cultures was obtained via filtration and drying to estimate fungal biomass in the inoculum. *S. pasteurii* (American Type Culture Collection #11859) starter cultures were started from frozen stock and grown in 100mL of BHI+ and incubated on a shaker table (New Brunswick Scientific, G24) at 30°C and 150RPM for 24 hours. Then, 1 mL of the *S. pasteurii* starter culture was pipetted into a new flask with fresh BHI+ and incubated (24h, 30°C, 150 RPM). After 24 hours, the sample was diluted with BHI to an optical density (OD) of 0.4 ± 0.02, or 2.13 ∗ 10^11^ ± 3.20^10^ CFU/mL of bacteria. This was accomplished by taking a 200µL aliquot from the sample and measuring OD at a wavelength of 600 nm using a plate reader (Biotek Synergy HT) and then diluting the sample accordingly. 1 mL of this diluted *S. pasteurii* culture was used to inoculate experimental samples.

### 5.3 Preparation of biomineralized mycelium scaffolds

To create self-mineralized fungal scaffolds, triplicate *N. crassa* cultures were started using the homogenized fungal inoculum (5.2) in 100 mL of either FICP-malt or FICP-def media. The cultures were covered from light and grown for 10 days on an orbital shaking table (New Brunswick Scientific, Innova 2300) at 23°C and 150 RPM. Liquid samples (60 µL) were taken on days 0, 1, 2, 3, 4, 5, and 10 for dissolved calcium and urea analysis. On day 10, fungal biomass from the cultures was removed via filtration using a glass fiber filter paper (Millipore Sigma AP1504700) and dried at 60°C until equilibrium dried mass was reached to measure dry weight of the mineralized fungal culture.

To bacterially mineralize mycelium scaffolds, fungal cultures were started from the inoculum protocol (2.2.1) and were then cultured in MBE media for 5 days without urea and calcium. These fungal cultures were autoclaved (121℃ and 20 psi) to kill the fungi. This was to ensure only bacteria were mineralizing the fungal scaffold.

*S. pasteurii* cultures were started using the protocol in 5.2. Autoclaved mycelium samples were rinsed in 10 mL of CMM-twice, and then placed into 250 mL Erlenmeyer flasks with 100 mL of CMM-medium. To start the growth of a bacteria on fungal scaffolds, 1 mL of *S. pasteurii* inoculum culture (2.2.2) was added into each flask with CMM- and autoclaved mycelium and incubated on a shaker table (New Brunswick Scientific, Innova 2300) for 24h at 23°C and 150 RPM. After 24 hours, *S. pasteurii* cultured with scaffolds were rinsed in CMM-twice and placed into new 250 mL Erlenmeyer flasks with 100 mL of CMM+ medium to induce mineralization and incubated (24h, 23°C, 150 RPM). Liquid samples (60 µL) were taken to analyze dissolved urea and dissolved calcium in solution at hours 0, 1, 4, 8, and 24. After 24 hours of cultivation, samples were dried at 60°C until equilibrium mass was reached.

To abiotically mineralize mycelium scaffolds (AICP), fungal scaffolds were first grown and autoclaved as described in 2.3.2. Autoclaved mycelium was then rinsed with DI water and placed into 50 mL of 11.098 g/L CaCl_2_ and water solution. 50 mL of 0.1 mM NaHCO_3_ and water solution was then added to the flask and mixed producing 100 mL of 0.05M CaCl_2_ and 0.05M NaHCO_3_ solution. The pH of the solution was raised to 8.5 using 40 g/L NaOH to induce calcium carbonate precipitation and was incubated on a shaker table (150RPM, 30 minutes, 23℃). Liquid samples (60 µL) were taken at minutes 0, 1, and 30 during the incubation for dissolved calcium assays.

### 5.4 Solution calcium and urea analysis

Two colorimetric assays were performed to measure dissolved calcium and dissolved urea present in the media throughout the biomineralization protocols. Biological triplicate cultures were sampled by diluting a 60uL sample from the culture into 540µL of 1.2M HNO_3_ to halt enzyme activity. Samples were stored at 4℃ before analysis. To determine changes in solution calcium concentrations throughout biomineralization, a modified calcium-o-cresolphthalein complexome method was used^3^. Briefly, 10 µL of diluted samples from an experiment were added to a 96-well plates with the assay reagents, shaken at 300 rpm for 1 minute, and incubated for 10 minutes at room temperature. The absorbance of samples within the 96-well plate was then measured using a Biotek Synergy HT plate reader at 575 nm.

Similar to the calcium assay, dissolved urea levels were measured throughout batch culture experiments to understand ureolysis occurring via *N. crassa* or *S. pasteurii*. Urea concentration was analyzed using a modified Jung assay^10, 62^. In the same manner as the calcium assay, 10 µL of diluted samples from an experiment were added to a 96 well plate with the assay reagents and shaken at 300 rpm for 1 minute. The well plate was then incubated at 30°C for 30 minutes. The absorbance of samples within the 96-well plate were then measured at 505 nm. Both the calcium and the Jung assays used technical triplicates and biological triplicates for each sample read with the plate reader.

### 5.5 Quantification of mineral content on biomineralized scaffolds

Calcium digests were performed in triplicate for FICP-malt, FICP-def, and BICP scaffolds to estimate the amount of biomineral precipitated onto mycelium scaffolds. Mineralized and non-mineralized mycelia were first dried at 60°C for 24 hours. Either 1.00g or 0.10g of mycelium, depending on available sample mass, was dissolved with 4 mL of 10% trace metal grade (TMG) nitric acid in 15 mL conical tubes for 24 hours at 22°C. The supernatant was then removed from the tubes. For samples with an ongoing visible reaction at 24 hours, digestion was continued for an additional 24 hours with another 10 mL of TMG nitric acid. Final dried mass was recorded after the digest, and after the sample had dried at 60°C for 24 hours. Final dried mass was normalized by the amount of sample added to the digest.

### 5.6 Characterization of biomineral composition and morphology

Portions of the dried mineralized scaffolds were sputter-coated with a thin, nanometer scale coating of iridium for scanning electron microscopy (Zeiss 55VP, 3kV accelerating voltage, working distance of 8.5-10 mm with an SE2 detector). Elemental mapping was performed with the same instrument and an electron dispersive spectroscopy (SEM-EDS) detector (Xplore 30 detector, Oxford Instruments) with a 12 kV accelerating voltage and a working distance of 8.5-10mm.

Another portion of the same dried samples were assessed for biomineral identities using a SCINTAG X1 X-ray Powder Diffraction Spectrometer (XRD). Approximately 1g per sample was crushed into a fine powder using a mortar and pestle and adhered to a glass slide with Vaseline and analyzed over a range of 2*θ* of 5–70. JADE (Materials Data) software was utilized for peak identification. Each biomineralized condition was assessed with biological triplicates.

### 5.7 Evaluation of biomineral microscale moduli

Portions from mineralized scaffolds were embedded in epoxy (PELCO epoxy, Ted Pella) and then ground to just below the samples surface with silicon carbide papers and then polished with a graded diamond suspension series to a final polish of 1 µm. Nanoindentation was performed on polished surfaces using a KLA-Tencor iMicro and a diamond Berkovich tip. Indents were performed using a trapezoidal load-hold-unload function of 30-60-30s to ensure viscoelastic energy had dissipated before unloading. Each indent had a maximum load of 3 mN with a 30 nm/s approach velocity. Arrays (10 × 10 indents with 20 *µ*m spacing in x and y) were placed on mineralized surfaces. Indents with load versus depth curves indicating strain hardening or excessive thermal drift were removed before data analysis. Indents optically confirmed to have been placed on epoxy or within 10 *µ*m of a mineral and epoxy interface were also excluded.

The reduced modulus (*Er*) was determined using Oliver-Pharr analysis ^63^. The contact stiffness, *S*, was calculated from the slop of the tangent line to a second-order polynomial fit to the 95th-45th percentiles of the unloading curve. *E_r_*, was calculated using *S* and the tip contact area, *A,* using Equation 1. *A* was calculated as a function of contact depth as determined from a calibration curve established on silica. The reduced modulus represents the composite elastic response of the tip (*E_t_*) and sample (*E_s_*). Tip properties are known (*E_t_* = 1140 GPa, *v_t_* = 0.07, but the sample Poisson’s ratio (*v_s_*) is not. *E_i_*, effectively a plain strain modulus, can therefore be calculated, which does not require assuming a Poisson’s ratio (**Equation 2**).

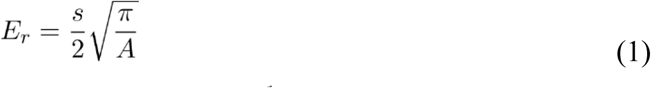

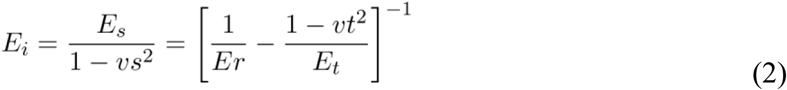

Following instrumented nanoindentation, epoxy embedded samples were further polished using a graded aluminum suspension to a 0.05 *µ*m finish. Atomic force microscopy (AFM, Asylum Research, Cypher S) was employed to acquire topography and modulus maps of biominerals. Given the high stiffness of calcium carbonate, a tip with a 200 N/m spring constant (RTESPA-525, Bruker) was chosen. The scan size was configured at 12 x 12 μm with a resolution of 384 x 384 pixels. To monitor potential tip wear or contamination, periodic calibration scans were performed on a glass surface with a known modulus (72 GPa, Fisherbrand); however, no significant tip wear was observed. Modulus maps were obtained using a trigger force of 1 μN and subsequently analyzed using a Hertzian model. Indents less than 5 GPa were excluded from the analysis to prevent curves placed on epoxy from influencing the data. Indents greater than 100GPa were also excluded as the stiffness of the cantilever does not allow for accurate measurements with moduli that high.

### 5.8 Visualization of bacterial growth on fungal scaffolds

To confirm bacterial attachment to the mycelium scaffold, confocal microscopy was performed on *S. pasteurii* that were cultured onto a mycelium scaffold. Fungal biomass was grown on a coupon to make handling the mycelium under the microscope simpler. Coupons were glass microscopy slides wrapped with a cellulose filter that was taped to remain attached to the slide. Coupons were then covered in aluminum foil and autoclaved. Coupons were inoculated with *N. crassa* and 100mL of MEB and incubated for 5 days at 150RPM and 23^°^C. *N. crassa* coupon cultures were autoclaved (Sec. 2.3.2). The flask containing the coupon with non-viable mycelium was then inoculated with *S. pasteurii* and cultured for 24 hours in CMM- at 150 RPM and 30^°^C (i.e., biocementation media but without calcium, to avoid mineralization obscuring bacterial attachment). Coupons were removed from the culture after 24 hours, rinsed with CMM- and placed into fresh CMM- and cultured for another 24 hours at 150RPM and 30^°^C.

Following the second incubation, the coupons was stained for imaging using propidium iodide and cyto-9 solution per manufacturer’s instructions (Thermo Fisher LIVE/DEAD BacLight L7007), removed from light to prevent bleaching, and incubated at 22℃ for15 minutes). The coupon was then rinsed with PBS. The mycelium was then stained using calcofluor white (Sigma-Adlrich 18909) a 0.25 g/L solution was prepared with Milli-q water, removed from any light source to prevent bleaching, and incubated at 22C, for15 minutes). The coupon was then rinsed again with PBS. A 1 cm^2^ sample of the scaffold was cut and removed from the center of the coupon and embedded in optimum cutting temperature fluid (Fisher Healthcare 4585) using dry ice^64^. These frozen scaffolds were cryosectioned (Leica CM1850l) into 20-µm slices at -20°C. Three serial slices were obtained from the top of the scaffold. Then, three 20-µm sections were sampled for every 500 µs until the bottom of the biofilm is reached. Finally, three serial slices were obtained from the bottom of the scaffold. Three biological replicates were obtained per condition. Confocal laser scanning microscopy of cryosectioned scaffolds was performed with a (Leica Stellaris 8 DIVE) using a 63x water objective. Z-stacks were taken with a 0.5-µ step size from the observed top of the cryosection to the observed bottom of the cryosection. Z-stacks were then analyzed using Imaris (Version 9.3.4 Oxford Instruments).

### 5.9 Determination of bacterial viability within biomineralized scaffolds

The viability of *S. pasteurii* on fungal mycelium scaffolds was tested at 2 and 4 weeks of drying and two different drying temperatures, room temperature (RT) or 30^°^C. Non-viable mycelium scaffolds were inoculated with *S. pasteurii* and either biomineralized (see section 2.3) or not mineralized (i.e., not provided mineralization media). Controls included fungal mycelium scaffolds without *S. pasteurii* and *S. pasteurii* planktonic cultures. At each time point (2 weeks and 4 weeks), two biological replicates of each drying condition (RT or 30^°^C) were fixed in a 2% glutaraldehyde and 4% paraformaldehyde solution and stored at 4°C for imaging. Three biological replicates were tested for viable *S. pasteurii*. Samples were incubated in 100 mL of BHI+ (150RPM, 23^°^C, 24h). Ureolytic activity was measured using a modified Jung assay at 0 hours and 24 hours, consistent with description in section 5.4. Afterwards, *S. pasteurii* viability was assessed from 1 mL aliquots using a bacterial plate count protocol.

To quantify viable bacteria, colony forming units (CFUs) were counted using a modified track-dilution protocol. 1 mL aliquots from liquid *S. pasteurii* cultures were diluted in series from 10^−1^ to 10^−8^ using PBS. 10 µL of each dilution was plated 5 times in a row on BHI agar supplemented with 20 g/L of urea. The inoculated agar plates were then incubated for 24 hours at 30^°^C. CFUs were counted for each of the 5 drops from the lowest dilution where CFUs were visible *CFU/mL* was then calculated using the following equation where *aCFU* is the average number of CFUs counted for 5 drops and *Df* is the dilution factor of the drops counted.

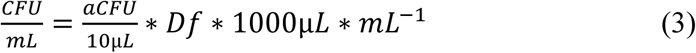

### 5.10 Determination of fungal viability within biomineralized scaffolds

The viability of mycelium scaffolds was estimated by their ability to grow on nutrient-rich agar plates. Growth measurements were made for biomineralized scaffolds (FICP-malt or FICP-def) and non-mineralized fungal scaffolds, after drying (room temperature or 30°C) for 2 weeks or 4 weeks. To determine the ability of scaffolds to grow after drying, scaffolds were placed on MBE agar plates, incubated (30°C, 5 days). If no growth was observed the scaffold was then ground into a paste using a sterilized mortar and pestle in an attempt to^65^ stimulate growth. The ground mycelium scaffold was then transferred to fresh MBE agar and incubated for 5 days at 30°C. Camera images of the MBE plates were analyzed using ImageJ to calculate the percent area covered by mycelium^66^. Measurements were completed for biological triplicates.

### 5.11 Construction of osteonal-mimetic microarchitectures using biomineralized mycelium

Osteonal-mimetic microarchitectures were constructed by growing *N. crassa* in a planar form, wrapping the planar mycelium scaffold around a plastic rod, and then mineralizing this mycelium scaffold with *S. pasteurii*. This was accomplished by inoculating 300mL of MBE in a baking dish with *N. crassa* (Sec. 5.2) and incubated for three days at room temperature without shaking. After incubation, the planar growth of *N. crassa* mycelium was autoclaved and rinsed with 50 mL of CMM- twice. The non-viable planar culture of *N. crassa* was added to a new baking dish with 300mL of CMM- and inoculated with 3mL of *S. pasteurii* inoculum (Sec. 5.2) and incubated at room temperature without shaking for 24 hours. Next, the planar scaffold with *S. pasteurii* was wrapped around a plastic rod and placed into a baking dish with 300mL of CMM+ media and incubated at room temperature without shaking for 24 hours to mineralize the wrapped scaffolds. After this final 24 hours, the mineralized, wrapped scaffolds, or artificial osteons, were cut into 1cm sections and dried at 60°C until constant mass was reached.

Prisms containing artificial osteons were manufactured by combining sand and artificial osteons into a PLA mold. Molds then undergo cyclic immersion in CMM- and CMM+ with *S. pasteurii*. The molds were rectangular prisms with interior dimensions of 2.54cm by 2.54cm by 10.16cm that were printed from a 3D printer. The print design included 1mm slits that ran along the height of each face to allow for fluid to flow throughout. Sand, prior to use, was soaked in 1.5M nitric acid for 24 hours, rinsed thrice with DI water, and finally autoclaved. Assembling the Osteonal-mimetic prisms began with filling a mold with treated sand to a third of its internal volume. Next a layer of the biomineralized mycelium scaffolds were packed, as tightly as possible, overtop the sand layer and covered in an additional layer of sand to the second third of the internal volume of the mold. Finally, another layer of tightly packed biomineralized mycelium scaffolds were added and covered with sand filling the mold. Molds were packed with just sand too to act as a control.

Molds were then mineralized. The packed molds are placed within a baking dish and immersed in CMM- (300mL per mold), making sure to not pour media directly on the molds and disrupt the loose sand within. The dishes with media and packed molds were inoculated with 3mL of 0.4 OD *S. pasteurii* culture (Sec. 5.2). The inoculated molds were incubated for 16 hours at room temperature with no shaking. After incubation the media was removed and was replaced with CMM+ (300mL per mold) and incubated for eight hours. This process is repeated six additional times for a total of seven culture and mineralization cycles. Following the final mineralization cycle the Osteonal-mimetic microarchitectures are de-molded, weighed, and loosely wrapped in aluminum foil before being placed in an oven at 60°C to dry until constant mass is reached.

## 6. Author Contributions

E.V: Conceptualization, Methodology, Formal Analysis, Investigation, Data Curation, Writing-Original Draft, Visualization, Writing-Review-Editing

E.H: Methodology, Investigation, Writing-Review-Editing

V.M: Methodology, Investigation, Writing-Review-Editing

S.L: Methodology, Investigation, Writing-Review-Editing

A.D: Software, Formal Analysis, Investigation, Writing-Review-Editing

L.M.C: Resources, Writing-Review-Editing, Supervision

A.P: Conceptualization, Writing-Review-Editing, Supervision, Funding Acquisition

R.G: Conceptualization, Writing-Review-Editing, Supervision, Funding Acquisition

E.E.O: Conceptualization, Resources, Writing-Original Draft, Writing-Review-Editing, Supervision, Funding Acquisition

C.M.H: Conceptualization, Formal Analysis, Resources, Data Curation, Writing-Original Draft, Writing-Review-Editing, Supervision, Project Administration, Funding Acquisition

## 7. Acknowledgements

The authors gratefully acknowledge support from the National Science Foundation (C.H., E.E.O., A.P., R.G.: 2036867; C.H., E.E.O, R.G., 2328351). Any opinions, findings, and conclusions, or recommendations expressed in this material are those of the authors and do not necessarily reflect the views of the National Science Foundation. The authors appreciate assistance from the staff of the Montana State University Center for Biofilm Engineering (Dr. Heidi Smith), the Image and Chemical Analysis Laboratory (Dr. Sara Zacher), and the Subzero Research Laboratory (Dr. Ladean McKittrick). Imaging was made possible by the Center for Biofilm Imaging Facility at Montana State University, which is supported by funding from the National Science Foundation MRI Program (2018562), the M. J. Murdock Charitable Trust (202016116) and the US Department of Defense (77369LSRIP). Additionally, the authors appreciate support and helpful conversations from the students within C.M.H, R.G, E.E.O, and A.P labs.

